# A common framework for semantic memory and semantic composition

**DOI:** 10.1101/2025.04.11.648390

**Authors:** Ryan M.C. Law, Matthew A. Lambon Ralph, Olaf Hauk

## Abstract

How the brain constructs meaning from individual words and phrases is a fundamental question for research in semantic cognition, language and their disorders. These two aspects of meaning are traditionally studied separately, resulting in two large, multi-method literatures, which we sought to bring together in this study. Not only would this address basic cognitive questions of how semantic cognition operates but also because, despite their distinct focuses, both literatures ascribe a critical role to the anterior temporal lobe (ATL) in each aspect of semantics. Given these considerations, we explored the notion that these systems rely on common underlying computational principles when activating conceptual semantic representations via single words, vs. building a coherent semantic representation across sequences of words. The present pre-registered study used magnetoencephalography and electroencephalography to track brain activity in participants reading nouns and adjective-noun phrases, whilst integrating conceptual variables from both literatures: the concreteness of nouns (e.g., “lettuce” vs. “fiction”) and the denotational semantics of adjectives (subsective vs. privative, e.g., “bad” vs. “fake”). Region-of-interest analyses show that bilateral ATLs responded more strongly to phrases at different timepoints, irrespective of concreteness. Decoding analyses on ATL signals further revealed a time-varying representational format for adjective semantics, whereas representations of noun concreteness were more stable and maintained for around 300 ms. Further, the neural representation of noun concreteness was modulated by the preceding adjectives: decoders learning concreteness signals in single words generalised better to subsective relative to privative phrases. These findings point to a unified ATL function for semantic memory and composition.

## 1. Introduction

Constructing meaning from individual words and sequences of words are fundamental challenges for research in semantic cognition, language and their disorders. These two aspects of semantics have formed separate research traditions, namely semantic memory and semantic composition. In the present study, we sought to unify these two literatures under a common framework. Not only would this effort address fundamental questions of how semantic cognition works, but also because, despite their distinct focuses, both literatures consistently implicate the anterior temporal lobe (ATL). Motivated by these two principal reasons, we explore the idea that activating conceptual semantic representations via single words, vs. building up meaning across sequences of words, might share underlying neurocomputational principles.

One perspective on the neurocomputational bases of semantic memory is the hub-and-spokes (H&S) model (for original model and reviews, see e.g., Rogers et al., 2004; Patterson et al., 2007; Lambon Ralph et al., 2017). ATL involvement in semantic memory is evidenced most notably in the syndrome of semantic dementia, in which atrophy to the ATLs leads to progressive deterioration in verbal and non-verbal conceptual knowledge (Patterson et al., 2007). On the other hand, semantic composition, or conceptual combination, has been investigated extensively in a minimal two-word phrase reading paradigm (for reviews, see e.g., Coutanche et al., 2019; Pylkkänen, 2019, 2020; Călinescu et al., 2023). These two large literatures have a striking convergence: they both implicate the left, and to a lesser extent the right, ATLs. Does the overlap in anatomical and functional localisation hint at shared computations across the two systems?

Seminal work by Pylkkänen and colleagues has begun to integrate these two research traditions in a series of MEG studies (Westerlund & Pylkkänen, 2014; Zhang & Pylkkänen, 2015; Ziegler & Pylkkänen, 2016; Zhang & Pylkkänen, 2018b; Kim & Pylkkänen, 2019; Pylkkänen & Brennan, 2020). Their work shows that the left ATL’s early involvement—200-250 ms post noun onset—in composition is modulated by conceptual specificity, suggesting that these two processes share common mechanisms at some level (Westerlund & Pylkkänen, 2014; Zhang & Pylkkänen, 2015; Kim & Pylkkänen, 2019). The present study builds on this work and aims to further explore in more detail what these underlying commonalities might be.

Formal, “open science” computationally-instantiated theories of semantic memory and sentence processing form a particularly relevant background for our purpose here, and we briefly overview each in turn. For semantic memory, the H&S approach models the ATL as a hidden ‘hub’ layer in a deep recurrent network, with reciprocal connections to ‘spokes’ layers each of which represents different object properties across sensorimotor and verbal modalities (Rogers et al., 2004; Lambon Ralph et al., 2010, 2017; Jackson et al., 2021). Input to any of these layers activates the corresponding pattern across the other layers (auto-associator) via the hidden hub layer, allowing for the extraction of transmodal and transtemporal statistical structures (which are the stable, coherent concepts) latent in verbal and non-verbal experiences across the lifespan. Crucially, the “hub” layer permits the representation of coherent concepts over and above linear feature combination and captures non-linear conceptual similarities that transcend superficial similarities (for an articulation of this idea in single concept explorations, see e.g. Lambon Ralph et al., 2010).

For semantic composition, computational models represent word meanings as vectors in high-dimensional spaces (Landauer & Dumais, 1997; Mikolov et al., 2013; Pennington et al., 2014) and implement composition as operations over constituent word vectors to derive phrase level representations, such as addition/multiplication (Mitchell & Lapata, 2008, 2010), tensor product (Smolensky, 1990; Clark & Pulman, 2007) (see also Martin & Doumas, 2019 for an alternative treatment to binding, a subroutine of composition). Other perspectives of sentence processing, such as the Sentence Gestalt models (McClelland et al., 1989; St. John & McClelland, 1990), conceive of sentence processing not as the building up of compositional structures. Instead, each word in a sentence serves as a cue to constrain and update a probabilistic representation about an event. This dynamic, context-dependent updating of representations also finds echoes in modern contextualized models like ELMo (Peters et al., 2018) and BERT (Devlin et al., 2019), where word meanings are not fixed but are derived based on their full sentence context.

A recent computational model by Hoffman et al. (2018) offers an explicit framework that converges models of semantic memory and semantic composition. The model takes a sequence of single words as input and learns by predicting both the upcoming words and each word’s sensory-motor properties (for the concrete words alone). Through this learning process, the model acquires context-sensitive semantic representations that reflect the multimodal experiential knowledge for each concept (as described in the standard H&S model) while also accounting for co-occurrence and information integration over time (McClelland et al., 1989; St. John & McClelland, 1990; Rabovsky et al., 2018). Indeed, because non-verbal experiences unfold over time, for concepts to be coherent and generalisable, this knowledge must be integrated both across modalities and across time (Jackson et al., 2021; Lambon Ralph et al., 2010). This integration is not a simple combination of features, but rather a nonlinear process crucial for capturing the complex relationships between different sensorimotor and verbal attributes of a concept. By assimilating evidence from these different strands, we propose that the ATLs provide a unified function essential to semantic representation—not just at longer timescales during learning but also during processing—and thus play a central role in both parallel literatures.

If there is indeed a unified transmodal, transtemporal integrated semantic “hub” function for the ATL, then adjective-noun processing should be sensitive to variables that are associated with individual concept strength which have been extensively studied in healthy and patient populations. One of the most prominent of these variables is concreteness or imageability. Concrete and abstract single nouns (e.g., *lettuce*/*fiction*) are processed differently, owing to either distinct representational formats (Paivio, 1971) or ease of information access (Schwanenflugel & Shoben, 1983) and yet are similarly supported by ventrolateral ATL (Hoffman et al., 2015). Importing this contrast into investigations of adjective-noun processing—which typically adopted concrete concepts—thus allows us to ask if neural systems supporting composition are generalised engines, processing input concepts with divergent formats and composing them alike.

To test our proposal that the same neurocomputational principles underlying semantic memory also underlie composition, we also integrated a different kind of semantic distinction of *denotation*, introduced by subsective vs. privative adjectives (e.g., *bad* vs. *fake*) (Kamp, 1975; Partee, 2010; Del Pinal, 2015). These adjectives transform conceptual representations in divergent and non-linear ways, yielding combined meanings that are not merely the sum of their parts—a challenge akin to computing coherent single concepts. For example, *bad* in *bad lettuce* does not just add a negative quality; it inhibits core *lettuce* properties (e.g., ‘is-green’) and activates properties that might be considered non-essential (e.g., ‘is-slimy’). Whereas, *fake lettuce* behaves rather differently viz. the properties it inhibits (e.g., ‘is-a-plant’) and promotes (e.g., ‘is-man-made’). These adjectives also introduce divergent *logical* semantics—a contrast less familiar to studies of semantic memory. The expression “this is a bad lettuce” entails “this is a lettuce”. Whereas, “this is a fake lettuce” does not entail “this is a lettuce”. A recent EEG study has begun exploring the processing profile of this contrast and found ERP effects around 500-800 ms post noun onset (Fritz & Baggio, 2020). Here, we asked if the ATL conceptual system is sensitive to this sort of logical semantics and explored how they transform conceptual representations.

In this pre-registered magnetoencephalography (MEG) and electroencephalography (EEG) study, we charted neural activity as 36 volunteers silently read nouns and adjective-noun phrases. Using mass-univariate region-of-interest analyses, we tested (1) whether neural systems supporting composition are generalised engines, composing concrete and abstract concepts alike, and (2) whether the ATL conceptual system is sensitive to logical semantics introduced by subsective-privative adjectives. Using multivariate decoding methods, we also tracked how ATL conceptual representations transform and evolve during composition.

## 2. Methods

The study was approved by the local ethics committee. The study was pre-registered on the Open Science Framework (OSF) on 14^th^ July 2023, before data collection began. The pre-registration document can be accessed on OSF at https://osf.io/m82nb/.

### 2.1 Participants

36 healthy adult participants (27 women; mean age = 26 years; SD = 6 years, range 18-40 years) participated in the study. The number of participants were calculated based on a power estimation (effect size *f* = 0.25, type I error rate α = 0.01, type II error rate β = 0.05) using *G*Power* (v3.1.9.6; Faul et al., 2007) on effect sizes reported in Bemis and Pylkkänen (2011). Participants were native English speakers, right-handed, without history of neurological disorder, had normal or corrected-to-normal vision, and were able to undergo magnetic resonance imaging (MRI).

Participants gave informed consent and were paid £12/hour for MEG and EEG, £12/hour for anatomical MRI scans, and £6/hour for stimulus norming (web-based sample separate from the MEG-EEG sample).

### 2.2 Stimuli

To create our stimuli, we selected nouns spanning a broad range of conceptual domains while controlling for key psycholinguistic properties. Using concreteness ratings from Brysbaert et al. (2014), we sampled words denoting concrete and abstract concepts. The rating norm contains human concreteness ratings range from 1 (fully abstract) to 5 (fully concrete). Concrete words were drawn from the range of 4.0-5.0, while abstract words were drawn from the range of 1.5-2.5; These ranges reflect the two local maxima in the distribution of mean concreteness ratings. From this pool, we used *LexOPS* (Taylor et al., 2020) to create concrete-abstract word pairs matched on variables known to influence lexical and conceptual processing, such as unigram Zipf frequency, age of acquisition, valence, and lexical decision latency (Table 1). All selected words were known by at least 90% of the raters, 3-10 letter long, noun-classified, and had low rating variability (SD < 1). We manually excluded items that were overly evocative or inappropriate, resulting in 137 matched pairs. These words span diverse conceptual categories, including objects, animals, food, sounds, places, emotions, and situations.

**Table 1.** Psycholinguistic characteristics of the final set of 100 nouns in each condition.

|  | Concrete | Abstract |
| --- | --- | --- |
| Concreteness rating | 4.55 (0.29) | 2.09 (0.27) |
| Imageability | 5.96 (0.66) | 3.28 (0.59) |
| Length (number of letters) | 5.80 (1.32) | 6.06 (1.24) |
| Zipf frequency | 4.11 (0.68) | 4.14 (0.67) |
| Age of acquisition (years) | 4.60 (2.82) | 5.49 (2.85) |
| Valence | 5.00 (1.79) | 5.08 (2.03) |
| Lexical decision latency (ms) | 570.02 (51.24) | 569.27 (46.58) |

*Mean values for each measure are reported, with SD in parentheses.*

We drew on insights from the field of linguistic theory and Natural Language Processing to inform our selection of adjectives. Privative adjectives were sourced from the list compiled by Nayak and colleagues (2014), while subsective adjectives were taken from Lalisse and Asudeh’s (2015) compilation. From each category, we selected 16 adjectives chosen for their broad compatibility with our noun set. To create the single-word condition, we generated length-matched consonant strings that served as visual placeholders lacking semantic content.

To create phrases, each noun was randomly paired with an adjective. As a coarse proxy of naturalness, we used Google Books Ngram data (Michel et al., 2011) to compute bigram frequencies, including dependency and sequence frequencies, for each phrase (Table 2). Dependency frequency captures instances where the adjective structurally modifies the noun, regardless of their proximity in the sentence, while sequential frequency captures instances where the adjective appears immediately before the noun. The bigram frequencies were summed within a set to compute aggregate ‘naturalness’ scores. Based on these aggregate scores, we selected the top 100 stimulus sets (600 items total; 100 per condition) for our final stimulus list. Table 3 shows an example set. For the full stimuli list, see https://osf.io/m82nb/.

**Table 2.** Zipf-transformed frequencies of each phrasal condition. Zipf scale values are computed as log₁₀(frequency per million) + 3. Frequencies are derived from Google Books (British English, 1919–2019).

|  | Concrete-Subsective | Concrete-Privative | Abstract-Subsective | Abstract-Privative |
| --- | --- | --- | --- | --- |
| Dependency frequency | -0.20 (1.64) | -0.15 (2.78) | 1.92 (4.08) | -0.69 (2.33) |
| Sequence frequency | -0.16 (1.72) | -0.18 (2.70) | 1.94 (4.10) | -0.75 (2.39) |

*Mean values for each measure are reported, with SD in parentheses.*

**Table 3.**
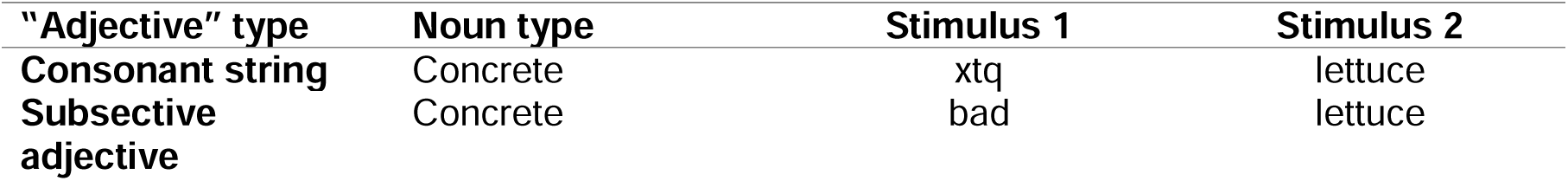
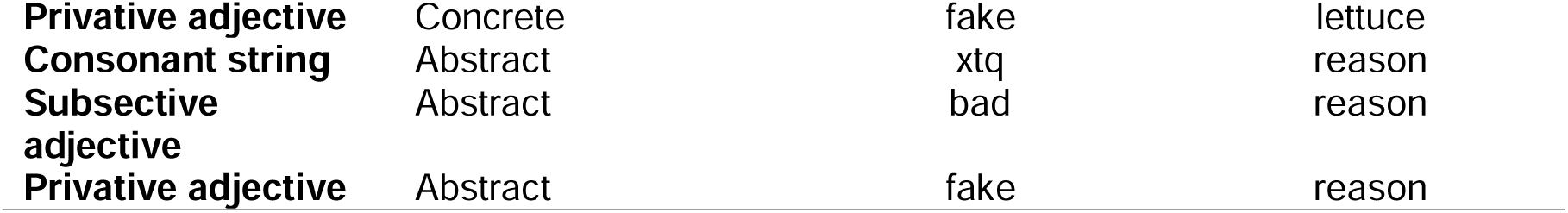
An illustrative set of items showing the study design.

To encourage participants to process the stimuli at the conceptual level, we opted for a phrase comprehension task (e.g. the stimulus “*bad lettuce”* followed by “*Is it crunchy?*”) instead of a picture-matching task as used in the semantic memory literature (e.g., Rogers et al., 2006) and the sentence processing literature (e.g., Bemis & Pylkkänen, 2011). This decision was based on two considerations: First, a picture-matching task is not suitable for abstract items. Second, following our manipulation of denotational semantics (subsective vs. privative), a comprehension task ensures that we probe compositional semantic processing, rather than simply aspects of individual adjectives or nouns. We created comprehension probes for 10% of trials (n=60). We note that the probe questions were designed to engage comprehension but were not matched for length or normed for difficulty. Consequently, the behavioural results reported here are largely descriptive and were not critical to the conclusions we drew.

### 2.3 Experiment procedure

Items were fully randomised and divided into five blocks (a brief break every 8 minutes or so). The stimuli appeared in light grey against black, backprojected onto a monitor about 90 cm away from the participant’s eyes. Each word or letter string subtended maximally at a horizontal visual angle of 4° and a vertical visual angle of .6°. Each trial began with a fixation cross in the centre of the screen for 300 ms, followed immediately by stimulus 1 (either letter strings or adjectives) on for 300 ms and off for 300 ms, stimulus 2 (noun) on for 300 ms and off for 500 ms. The decision to allow for a longer period post second word onset (800 ms) is based on Fritz and Baggio’s (2020) finding that denotation effects emerge late (500-800 ms). After 10% of trials there was a yes/no comprehension question, presented in whole and in cyan, prompting a right-hand button press response: index finger for yes and middle finger for no. A further inter-trial interval of 1.5 s on average (uniformly jittered between 0.75-1.25 s) was included for cortical activations to return to baseline (Hansen et al., 2010) (Figure 1). The whole experiment inside the MEG chamber took around 45 minutes (excluding EEG preparation time, instructions, etc.).

**Figure 1.**
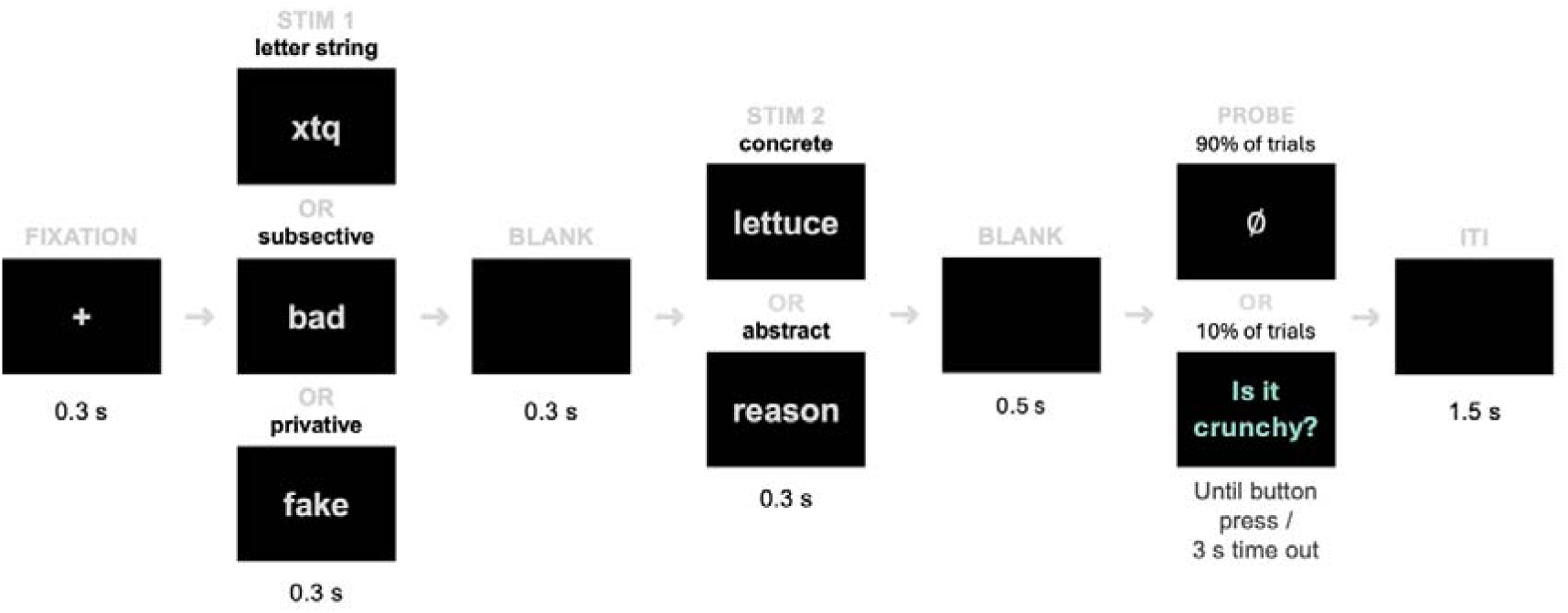
Trial structure. Task probes were included for 10% of trials to encourage attention.

### 2.4 MEG-EEG data acquisition and preprocessing

We collected concurrent MEG and EEG recordings to maximize spatial resolution (Molins et al., 2008) as well as structural MR images for each participant. MEG and EEG data were acquired in a magnetically shielded room using a 306-channel TRIUX neo system (MEGIN, Espoo, Finland), with 204 planar gradiometers and 102 magnetometers. EEG data were collected concurrently using a 64-electrode EEG cap (EasyCap GmbH, Herrsching, Germany) with a 10-20 layout. EEG reference and ground electrodes were attached to the nose and left cheek, respectively. The electrooculogram (EOG) was recorded by electrodes above and below the left eye (vertical EOG) and at the outer canthi (horizontal EOG), respectively. Impedance levels were reduced to 10 kOhms or lower during preparation when possible. Prior to data acquisition, the positions of 5 head position indicator coils attached to the EEG cap, 3 anatomical landmark points (two preauricular points and nasion) as well as approximately 50–100 additional points covering the scalp were digitized using a 3Space Isotrak II System (Polhemus, Colchester, Vermont, USA) for later co-registration with MRI data. EEG and MEG data were digitized at a sampling rate of 1 kHz and with a high-pass filter of 0.03 Hz. As for MR images, for each participant, a T1-weighted structural scan was acquired using a Siemens 3T Prisma scanner.

We then used the Python package *MNE-Python* (v1.4.2; Gramfort et al., 2013, 2014) to pre-process the MEG and EEG data. Data were bandpass-filtered offline between 0.1-40 Hz with a finite impulse response filter design using a Hamming window method. Data were subject to the following noise reduction procedures. For MEG data, we first identified and repaired bad and noisy channels. We then applied Signal Space Separation (SSS) with its spatiotemporal extensions to reduce environmental artefacts (Taulu & Kajola, 2005). This step also included head movement compensation. For EEG, bad channels were visually identified and interpolated using the spherical spline method (Perrin et al., 1989). To attenuate eye-movement artefacts, we performed an independent component analysis (ICA) using the *FastICA* algorithm (Hyvärinen & Oja, 2000). Eye-movement related components were detected using correlation with EOG channels and were inspected visually and removed. Filtered and noise-reduced continuous data were then segmented into epochs from 0.2 s before fixation cross onset to 1.4 s post first word onset. Baseline correction was applied using signals spanning 0.2 before fixation cross onset. We applied a common baseline for all epochs. Epoched data were then downsampled to 250 Hz.

Epochs were first rejected using an automated procedure using the Python package *autoreject* (Jas et al., 2017). All trials were included in further analysis since the comprehension trials are to encourage attention and comprehension. This procedure yielded on average around 80 trials per condition for further analyses.

### 2.5 Source reconstruction

We reconstructed distributed cortical source estimates by combining MEG and EEG within a minimum-norm estimation (MNE) framework (Hämäläinen & Ilmoniemi, 1994; Hauk, 2004; Hauk et al., 2022). This yielded a time-series of cortical current dipole moment at each vertex in a common source space. Below, we describe each step in turn, from anatomical reconstruction to source estimation.

Each participant’s T1-weighted structural MRI was first segmented to reconstruct subject-specific anatomical models of the scalp, inner and outer skulls, and cortical surface using the automated segmentation algorithms (Dale et al., 1999) in *FreeSurfer* (v6.0.0; Fischl, 2012). Using *MNE-Python* (v1.4.2; Gramfort et al., 2013, 2014), we constructed a three-layer boundary element (BEM) model using those surfaces (conductivity parameters are 0.3, 0.006, and 0.3 respectively). The MEG-EEG sensor configurations and MR structural images were then co-registered to the reconstructed scalp surface by aligning fiducial landmarks (nasion, left and right preauricular points) and approximately 50-100 additional head-shape points.

Next, we defined a source space consisting of4098 potential dipole locations on each hemisphere’s surface (8,196 dipole locations in total), with three orthogonal orientation components at each vertex. To calculate the inverse solution, we estimated the noise covariance matrix using the 200 ms window prior to fixation and then regularised it using an automated model selection method (Engemann & Gramfort, 2015). MEG and EEG data were then “whitened” to ensure that different channel types contribute to source reconstruction appropriately with respect to their differing noise levels or units (MEG measured in fT, EEG in µV). The BEM and source space were used to compute the forward solution.

The regularised noise covariance matrices and forward solutions were used to assemble the inverse operator using L2-MNE with a loose orientation constraint value of 0.2 and depth weighting of 0.8. The inverse operator was then regularised assuming a signal-to-noise ratio (SNR) of 3. When applying the inverse operator to the combined MEG–EEG sensor data, the output at each cortical dipole is expressed as a dipole moment (Am). After applying the inverse operator to sensor evoked responses, individual-specific cortex-wide source estimates were morphed to a common space (‘*fsaverage’*) for group-level mass-univariate analyses.

For our decoding analyses, we reconstructed source estimates with a near-identical pipeline, with only one point of divergence: that the inverse operator was assembled with a fixed orientation constraint (i.e., normal to the cortical surface), yielding signed data that retains directionality of the current flow which may help boost decoder performance.

### 2.6 Analyses

#### 2.6.1 Mass-univariate region-of-interest (ROI) analyses

Our investigation primarily targeted the left and right ATLs. Bilateral ATLs are strongly implicated in semantic memory (e.g., Rogers et al., 2006; Patterson et al., 2007; Lambon Ralph et al., 2017), and while the left, and to a lesser extent the right, ATL, are implicated in semantic composition (Pylkkänen, 2019, 2020). Further analyses also included secondary ROIs, namely the IFC and posterior-lateral temporal lobe (PTL), which have been implicated in studies of semantic control (Jackson et al., 2021, 2021; Lambon Ralph et al., 2017). This inclusion was motivated by theoretical work that posit additional processing (e.g., “conceptual blending”, Coulson & Fauconnier, 1999; “coercion”, Partee, 2010) for privative expressions. The results for these secondary ROIs are reported in the Supplementary Materials.

ROIs were defined based on four MNI coordinates (Table 4) on the cortical surface. We created labels with a radius of 30-mm (40-mm for the ATLs) around each coordinate (Figure 2) (see Supplementary Figure S1 for a visualization of all ROIs). These labels were chosen to include key brain activations observed in previous work on phrase comprehension (Bemis & Pylkkänen, 2011; Pylkkänen, 2019, 2020) and functional MRI (Pallier et al., 2011) and semantic cognition, including meta-analyses (Lambon Ralph et al., 2017; Jackson, 2021). To include the corresponding right hemisphere regions, we reversed the x-coordinate of each label.

**Figure 2.**
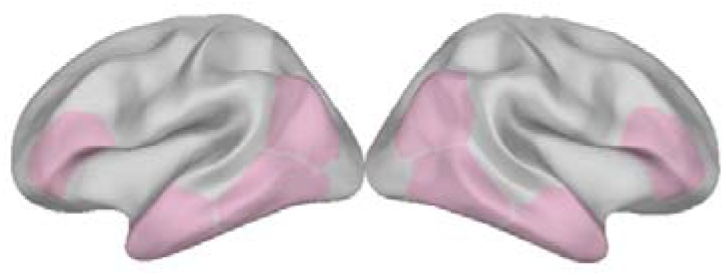
Regions of interest.

**Table 4.** MNI coordinates of ROI seeds for growing ROI labels.

| ROIs | MNI coordinates | ROIs | MNI coordinates |
| --- | --- | --- | --- |
| Left ATL | [-57, -2, -33] | Right ATL | [57, -2, -33] |
| Left PTL | [-50, -52, -6] | Right PTL | [50, -52, -6] |
| Left IFC | [-45, -66, 24] | Right IFC | [45, -66, 24] |
| Left TPC | [-51, 30, 6] | Right TPC | [51, 30, 6] |

We used mass-univariate mixed-effects ANOVAs combined with non-parametric cluster-based permutation tests (Maris & Oostenveld, 2007) to analyse brain activity in each ROI over time. This was done using the Python package *Eelbrain* (version 0.39; Brodbeck et al., 2023). We first averaged the cortical activity within each ROI to get a single time course for each. Then, to address our specific research questions, we fitted different statistical models at each time point for each ROI, analysing the full 800 ms time window spanning the second word.

The first aim of the study was to better understand the generality of the ATLs in composing concrete as well as abstract concepts. Thus, the following analyses were performed for the ATL first. However, as mentioned above, privative expressions (e.g., *fake lettuce*) may incur additional processing costs (Coulson & Fauconnier, 1999; Partee, 2010). Prior work on semantic memory also reported a “concreteness advantage” (Hoffman, 2016) in which the comprehension of abstract words additional recruit semantic control regions (Hoffman et al., 2015). Therefore, we also separately tested secondary ROIs, including IFC and PTL, as well as TPC.

We first tested for previously reported effects of Composition (single word, subsective phrase) and Concreteness (concrete, abstract), and also included a factor Hemisphere (left, right) in ATLs. To this end, we fitted a 2x2x2 mixed-effects ANOVA at each time point separately, with three fixed-effects factors Composition, Concreteness and Hemisphere. One major motivation to include hemisphere as a factor was that both left and right ATLs contribute to conceptualisation (Lambon Ralph et al., 2017). If the ATLs are generalised engines, composing concrete and abstract concepts alike, we would expect a main effect of Composition (subsective phrase>word), with engagement for both concrete and abstract meanings.

A second aim of the study was to test whether the ATL conceptual system is sensitive to logical semantics introduced by subsective and privative adjectives). Previous MEG studies targeting the left ATL found that it is sensitive to logical semantics during phrase processing (Zhang & Pylkkänen, 2018b, 2018a). Given this background, to study the encoding of logical semantics in the left ATL, we used a 2x2 mixed-effects ANOVA separately at each time point, with the fixed-effects factors of Denotation and Concreteness. Recent EEG work found more negative-going potentials for privative relative to subsective phrases, (Fritz & Baggio, 2020), but we did not have strong predictions with respect to directionality of the effect in unsigned source-localised ATL signals.

For our secondary ROIs, we tested for effects of Composition, Concreteness, and Denotation, using a 2x2 mixed-effects ANOVA separately at each time point, with two fixed-effects factors Concreteness and Composition, and a 2x2 mixed-effects ANOVA separately at each time point, with factors Concreteness and Denotation.

Finally, we sought to further characterise the spatial specificity of any significant effects by testing for interaction effects with the factor, Region. For example, in order to follow up composition effects, the interaction analyses would consist of 2x2x4 mixed-effects ANOVAs with factors Concreteness, Composition, and Region (ATL, PTL, IFC, TPC).

In all models, subjects were included as a random effect.

To quantify the magnitude of the observed effects, we calculated the Cohen’s *f* as a measure of effect size (Cohen, 2013) for temporal clusters associated with significant effect. Cohen’s *f* is calculated as the standard deviation of the condition means divided by the common population standard deviation. We estimated the statistical significance of temporal clusters using cluster-based permutation tests per ROI in the time domain, with 5,000 randomisations. Since we ran a separate cluster-based permutation test for each ROI, this resulted in eight statistical tests. To correct for multiple comparisons across ROIs, we applied a False Discovery Rate correction using the Benjamini-Hochberg procedure (Benjamini & Hochberg, 1995) and reported corrected *q*-values (i.e. adjusted p-values).

To quantify the evidence for the presence or absence of an effect, we performed a Bayes factor analysis using the *Pingouin* package in Python (Vallat, 2018). The Bayes factor (BF10) expresses how many times more likely the data are to have occurred under the alternative hypothesis (H1) than the null hypothesis (H0). For instance, a BF10 of 3 indicates the data are three times more likely under H1 than H0.

Following conventions (e.g., Wagenmakers et al., 2018), we interpret BF10 values greater than 3, 10, 30, and 100 as ‘moderate’, ‘strong’, ‘very strong’, and ‘extreme’ evidence in favour of the alternative hypothesis, respectively (e.g., that neural activity from phrases was stronger than that from words). Conversely, the inverse values (BF10 < 1/3, 1/10, 1/30, and 1/100) provide corresponding levels of evidence in favour of the null hypothesis (e.g., that neural activity from concrete items was identical to that from abstract items).

#### 2.6.2 Multivariate decoding analyses

To determine if ATL brain signals encode information about a particular experimental condition (like whether a word is concrete or abstract), our analysis used temporal decoders. These decoders were trained at each time point to predict the condition. Their performance was measured over time using the area under the curve (AUC) score, creating a time course that reveals when the brain activity encodes information about the condition. We also performed temporal generalisation analyses (King & Dehaene, 2014), which involves training a decoder at one time point and testing its performance on all other time points. This creates a matrix that shows not only how well the decoder performs at any given moment, but also whether the neural patterns remain stable over time. In this matrix, the main diagonal represents the decoder’s performance at each time point, while the off-diagonal entries indicate how consistent the neural signals are across different points in time.

We used source-localised signals from the left ATL as input for our decoding models. To improve the SNR of the data, we created “pseudo-trials” by averaging several trials together, depending on the specific contrasts we were interested in: For subsective/privative adjective semantics, we averaged pairs of trials across the two levels of concreteness (80 trials each) within each denotation level, resulting in around 80 trials per adjective type. For concrete/abstract noun semantics, we averaged three trials across the three levels of adjective types (80 trials each) within each concreteness level, resulting in around 80 trials per noun type. Finally, we conducted separate temporal decoding analyses. For each time point, we trained and tested a unique decoder to distinguish between either concrete and abstract trials or subsective and privative trials.

To address the question of how adjective semantics shape noun representations, we trained decoders to distinguish concrete vs. abstract in single words and tested their ability to generalise separately to subsective and privative phrasal contexts. For this, we also similarly performed averaging separately within each test condition, yielding approximately 40 trials per condition. Similar to above, we performed a temporal decoding analysis, fitting a decoder at each time point to distinguish between concrete vs. abstract trials. However, recent EEG work exploring similar contrasts in univariate ERP responses informed our analysis (Fritz & Baggio, 2020). The authors explored denotational aspects of meaning and identified a cluster around 500-800 ms post noun onset. We expected that, in our analysis, the impact of our adjective manipulation would result in differential decodability similarly later, but not early, in neural activity. We took the cluster extent that Fritz and Baggio (2020) found to define a “late” time window, spanning 500-800 ms). As for an “early” time window, we drew insights from the well-characterised N400 ERP component and its relevance in meaning processing (Kutas & Federmeier, 2011) and defined a 200 ms window from 300-500 ms (See Figure 6). To statistically test for interactions between phrase type (subsective vs. privative) and time window (early vs. late), we fitted a linear mixed effects model to averaged neural responses within each time window using the *lme4* package in R (Bates, Mächler, Bolker, & Walker, 2015). The model included, as fixed effects: phrase type (subsective, privative), time window (early, late), an interaction term (phrase type by time window). The random-effects structure was initially specified as a maximal model (Barr et al., 2013), including by-subject random slopes and intercepts for both main effects and their interaction, and was then iteratively simplified to avoid singular fit or non-convergence. The final model included by-participant random intercepts and correlated slopes for time window and phrase type. *p*-values were computed using the *lmerTest* package (Kuznetsova et al., 2017). To estimate model fit, we computed marginal and conditional R² (Nakagawa & Schielzeth, 2013) using the *MuMIn* package (Bartoń, 2025). Follow-up post-hoc pairwise comparisons for significant effects were conducted using the *R* package *emmeans* (Lenth, 2024), and p-values adjusted using Tukey’s method as needed.

To prepare our data for decoding, we first standardised the time series from each ROI using the RobustScaler from *scikit-learn* (Pedregosa et al., 2011, 2011). The Scaler removes the median and scales the data using its interquartile range, as this method is more robust to outliers compared to the approach of removing the mean and scaling to unit variance. All decoding was performed within-subjects using a stratified 10-fold cross-validation. Decoders were L2-regularised logistic regression models from *scikit-learn* with default settings. For each subject, decoding performance was summarised with an area under the curve (AUC) score for each decoder for each fold then averaged across fold. Finally, we used one-tailed cluster-based permutation tests to compare these average scores against a chance level of 0.5 to determine group-level statistical significance.

#### 2.6.3 Behavioural analyses

As aforementioned, comprehension probes engaged comprehension and attention but were not matched for length or normed for difficulty, so behavioural data were reported for completeness and were not central to our conclusions.

To analyse reaction times (RT) data, we excluded incorrect trials. We fitted a linear mixed effects regression model to group-level log-transformed RT data using the *lme4* package in R (Bates, Mächler, Bolker, & Walker, 2015). The model included fixed effects for concreteness (concrete, abstract), denotation (baseline, subsective, privative), and an interaction term (concreteness by denotation). To analyse response accuracy (this time incorrect responses included), we fitted a generalised mixed effects logistic regression with a binomial distribution.

The random-effects structures were initially specified as a maximal model (Barr et al., 2013) for both models, including by-item random intercepts and by-subject random slopes and intercepts for both main effects and their interaction, and was then iteratively simplified to avoid singular fit or non-convergence. The final models for both accuracy and RT included by-participant and by-item random intercepts. *p*-values were computed using the *lmerTest* package (Kuznetsova et al., 2017). To estimate model fit, we computed marginal and conditional R² (Nakagawa & Schielzeth, 2013) using the *MuMIn* package (Bartoń, 2025). Follow-up post-hoc pairwise comparisons for significant effects were conducted (on data back-transformed from log scale to original scale) using the *R* package *emmeans* (Lenth, 2024), and *p*-values adjusted using Tukey’s method as needed.

## 3. Results

### 3.1 Behavioural results

For accuracy, a mixed-effects model found a main effect of denotation (χ² = 22.49, p < .0001) and a significant interaction between denotation and concreteness (χ² = 9.18, p = .010). There was no main effect of concreteness (χ² = 2.85, p = .091). The marginal R² was 0.12 and conditional R² was 0.32. Post-hoc pairwise comparisons on estimated marginal means revealed that accuracy was significantly lower for privative phrases relative to single-word controls (odds ratio = -2.61, *p* = .004) and to subsective phrases (odds ratio = -4.48, *p* < .0001). No significant difference was found between baseline and subsective phrases (odds ratio = 0.58, *p* = .20). Follow-up analyses on the concreteness-by-denotation interaction revealed that it was driven by an accuracy advantage for concrete subsective phrases relative to other conditions. This condition was significantly more accurate than both its abstract counterpart (log odds = 1.53, *p* = .0008) and other concrete conditions, i.e. concrete single-words (log odds = 1.19, *p* = .028) and concrete privative phrases (log odds = 2.42, *p* < .0001).

As for reaction times, we found main effects for concreteness (χ^2^ = 17.84, *p* <.0001) and denotation (χ^2^ = 21.47, *p* < .0001) but no interactions (χ^2^ = 3.91, *p* = .14). The marginal R² was 0.11 and conditional R² was 0.54. Across denotation types, concrete phrases were responded to more quickly (M = 1.52 s, SE = 0.06) than abstract phrases (M = 1.74 s, SE = 0.07), consistent with a ‘concreteness’ advantage (Hoffman, 2016). RTs also varied by denotation type. Participants were slowest to respond to privative phrases (M = 1.79 s, SE = 0.07), intermediate for baseline nouns (M = 1.60 s, SE = 0.07), and fastest for subsective phrases (M = 1.50 s, SE = 0.06). Post-hoc pairwise comparisons revealed participants were significantly faster for single-word baselines compared to privative phrases (ratio = 0.894, SE = 0.035, *t*(54.8) = −2.86, p = .016). No significant difference was observed between baseline and subsective phrases (ratio = 1.069, SE = 0.041, *t*(52.0) = 1.73, p = .205). Privative phrases were responded to significantly more slowly than subsective ones (ratio = 1.196, SE = 0.047, *t*(54.2) = 4.57, p < .001).

In all, these results are compatible with prior findings (1) of a “concreteness advantage” (Hoffman, 2016) and that (2) privative semantics incur additional processing costs, in line with predictions from linguistic theory (Coulson & Fauconnier, 1999; Partee, 2010) and empirical findings (Fritz & Baggio, 2020). We note again that the comprehension probes used here were not normed for difficulty but serve to promote conceptual processing. Thus, our central claims are not based on our behavioural data.

### 3.2 The ATLs are generalised engines for semantic composition

In our three-way ANOVAs of ATL activity with factors Composition, Concreteness and Hemisphere, we observed main effects of Composition (smallest *p* = .002) in the left and right ATL (Figure 3A). Composition effects were driven by temporal clusters around 390-580 ms (Cohen’s *f* = 0.09) and 590-660 ms (Cohen’s *f* = 0.08) post-noun-onset. In both cases, subsective phrases elicited higher activity relative to single-word controls (BF10 = 11.58, strong evidence; BF01 = 0.28). See Supplementary Figure 1 for the time courses of all tested ROIs (including secondary ROIs; see Methods) and Supplementary Figure 2 for uncorrected spatiotemporal t-maps. We also observed a main effect of Hemisphere (*p* = .0002), with stronger activity in the left than right ATL (Figure 3B).

**Figure 3.**
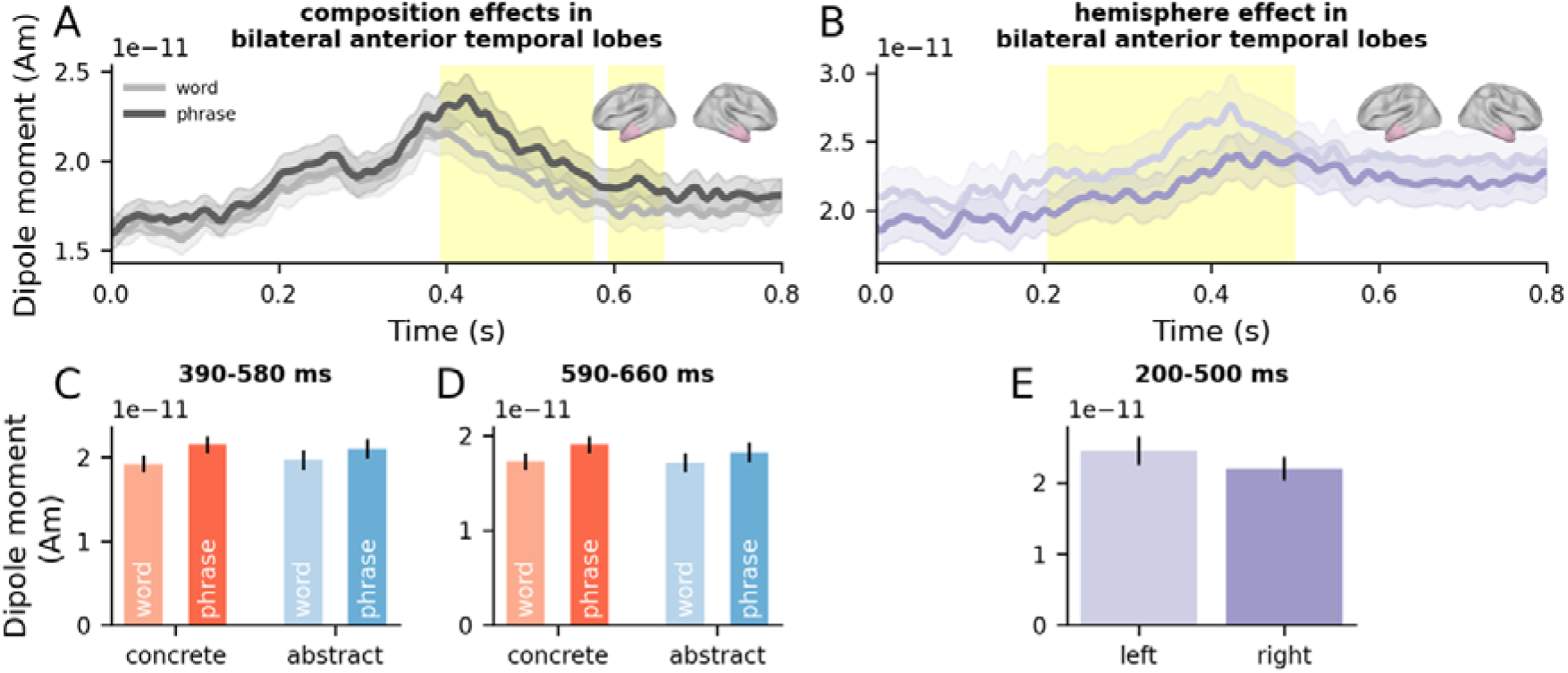
Time-resolved analyses of the ATL ROIs. (A, B) Time series of evoked responses in the left and right ATL, respectively, for words and subsective phrases, aligned to noun onset (0 s). Error bars represent ±1 within-subjects SEM (Loftus & Masson, 1994). Cluster extent identified from permutation testing are indicated by yellow shaded areas for significant effects (p<.05) and grey shaded areas for marginally significant effects (p<.1). (C, D, E) Mean response within the cluster time window, plotted separately for each condition. Error bars represent ±1 within-subjects SEM (Loftus & Masson, 1994). Units: Am = Ampere-meter, S = second.

Conversely, no Concreteness effects were found in the ATLs (*all p-values* > .05). Bayesian analysis further indicated moderate evidence in favour of the null hypothesis in the ATLs (BF10 = 0.34, BF01 = 3.15). See Supplementary Figure 5 for the time course of Bayes Factor evidence for Composition and Concreteness in these regions.

To further characterise the spatial specificity of these effects, we also examined Composition effects in all individual regions including secondary ROIs (see Methods) and again found Composition effects only in the left and right ATLs.

### 3.3 Left ATL is sensitive to adjectival types associated with phrasal interpretation

The statistical evidence for a Denotation effect in the left ATL was mixed. The cluster-based permutation analysis yielded a borderline effect (*p* = .052). However, a subsequent Bayesian analysis provided moderate evidence for the alternative hypothesis (BF10 = 3.69, BF01 = 0.39). The associated cluster spanned 380-440 ms post-noun-onset (Figure 4), characterised by increased activity for subsective relative to privative phrases (Cohen’s *f* = 0.08). See Supplementary Figure 3 for the time courses of all tested ROIs (including secondary ROIs; see Methods) and Supplementary Figure 4 for uncorrected spatiotemporal t-maps.

**Figure 4.**
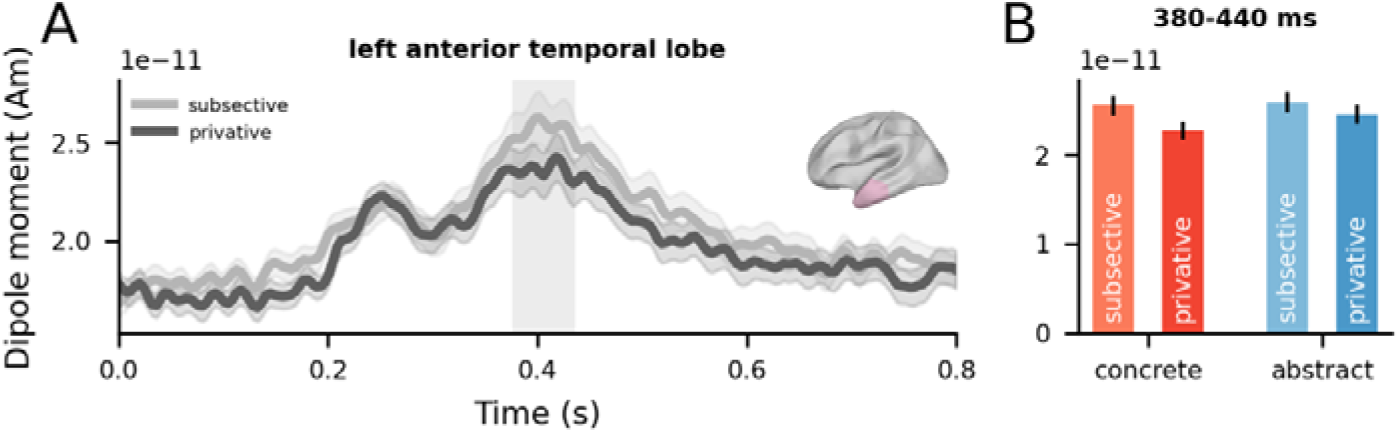
Time-resolved analyses of the left ATL ROI. (A) Time series of evoked responses in the left ATL for subsective and privative phrases, aligned to noun onset (0 s). Error bars represent ±1 within-subjects SEM (Loftus & Masson, 1994). Permutation-based clusters are indicated by grey shaded areas for marginally significant effects (p<.1). (B) Mean response within the cluster time window, plotted separately for each condition. Error bars represent ±1 within-subjects SEM (Loftus & Masson, 1994). Units: Am = Ampere-meter, S = second.

No significant effects of Concreteness were found, and neither did Concreteness modulate effects of Denotation. A Bayes Factor analysis further indicated weak evidence in favour of the null hypothesis in the left ATL (BF10 = 0.51, BF01 = 2.66). See Supplementary Figure 6 for Bayes factors time series for the effect.

### 3.4 Distinct representational formats for adjective and noun semantics

To characterise how individual adjective and noun semantics are represented, we trained decoders on neural signals from the left ATL at each time-point separately and tested those decoders on held-out data at those time points. Cluster-based permutation tests revealed above-chance decoding for adjective and noun semantics (all *p*-values < .05). Cluster extents associated are 290-360 ms, 410-480 ms, 550-610 ms and 620-680 ms for denotation (Figure 5A) and 280-490 ms and 500-560 ms for concreteness (Figure 5C).

**Figure 5.**
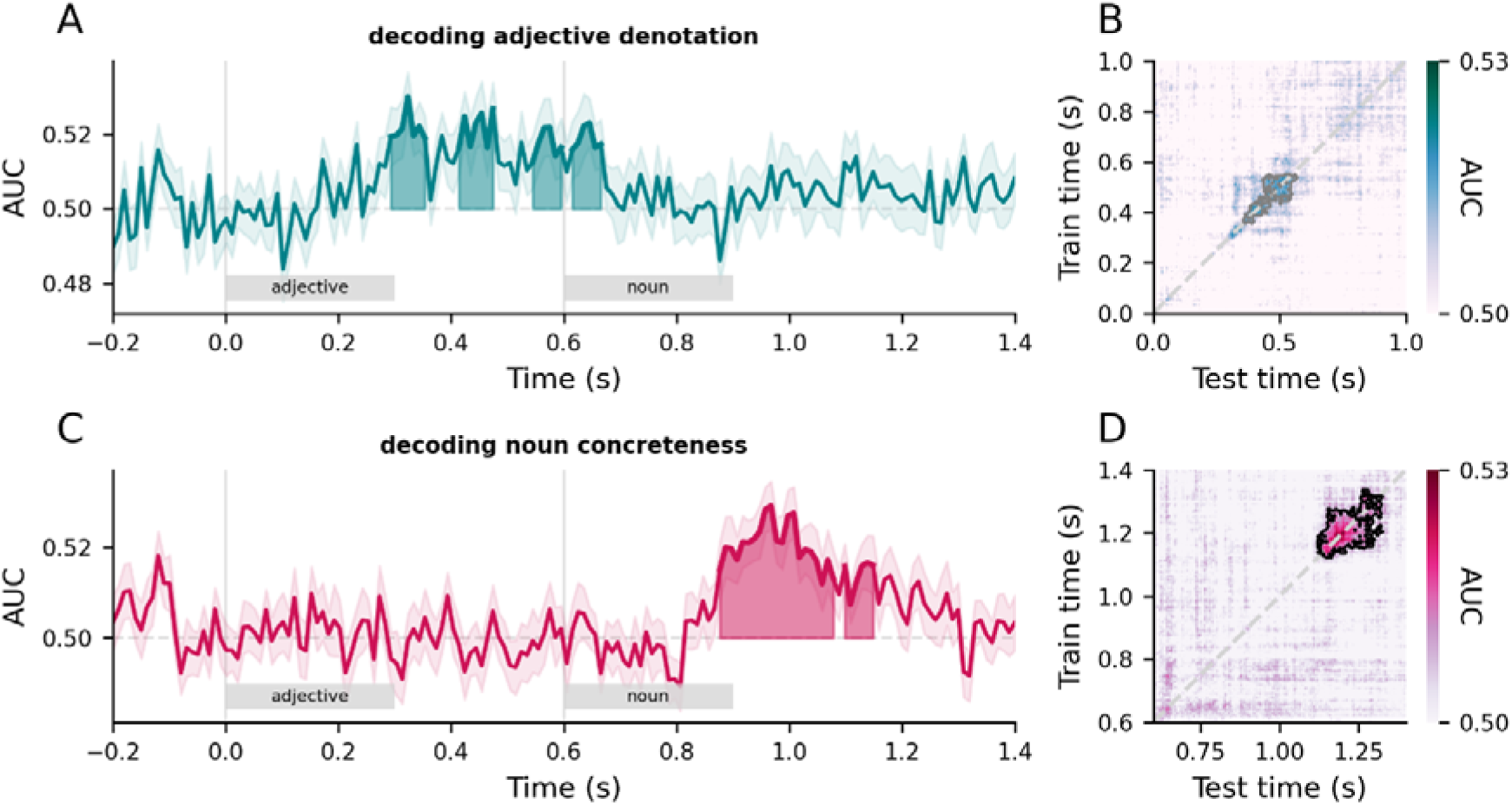
(A, C) Time series of decoding performance (AUC: Area under the curve summarising decoding performance) in the left ATL for adjective (A) and noun (C) semantics, aligned to adjective onset at 0 s. Error bands represent ±1 SEM. Shaded regions indicate cluster extents corresponding to group-level effects assessed with cluster-based one-sample permutation tests against chance. (B, D) Temporal generalisation matrices (TGM) of adjective (B) and noun (D) semantics, where time 0 s is adjective onset and 0.6 s is noun onset. Black and grey outlines denote cluster extents corresponding to significant and marginally significant group-level effects assessed with cluster-based one-sample permutation tests against chance.

We then used temporal generalisation to test whether the representations of adjective and noun semantics are maintained in a stable format during phrase comprehension. Cluster-based permutation tests revealed modest above-chance decoding for adjective (*p* = .088; Figure 5B) and robust above-chance decoding for noun semantics (*p* = .028; Figure 5D). To explore the time stability of the format, we calculated how far from the diagonal a given cluster extends by calculating the mean and max offset distances from the diagonal in the given cluster. The denotation cluster subtended a mean offset from the diagonal of ∼45 ms and a max offset of ∼150 ms. Whereas, the cluster associated with concreteness decoding was nearly twice as stable in time, with a mean offset from the diagonal of ∼80 ms and a max offset of ∼270 ms.

These results imply that the brain encodes the distinction between subsective and privative semantics at time points between 300-700 ms post adjective onset but the neural patterns changed from time point to time point. In contrast, the format of the representation of concrete vs. abstract semantics appeared to be more stable across time, characterised by longer off-diagonal decodability. For example, decoders trained at 1.1 s were able to generalise up to 1.3 s and vice versa, implying a recurring, stable format for representing conceptual semantics.

### 3.5 Adjective semantics differential impact noun representation

After characterising the temporal representation of these concepts, we sought to understand how noun representations evolve as a function of adjective semantics. In this analysis, decoders were trained on single word trials, distinguishing between concrete and abstract items. The decoders were then evaluated on how well they generalised to subsective and on privative trials separately. Cluster-based permutation tests on the decoding scores from the entire noun time window revealed above chance decoding in subsective phrases (*p* = .003; Figure 6A) and not in privative phrases (Figure 6C). The cluster extended from 330 to 440 ms post noun onset. We followed up with linear mixed effects modelling to explore the interactions between phrase type (subsective vs. privative) and time window (early vs. late) and found a main effect of time window (*p* = .021) and a main effect of phrase type (*p* = .037; Figure 6E) and no interaction (*p* = .820). Post hoc comparisons confirmed that decodability was higher in the early compared to the late time window (*t*(34) = 2.42, *p* = .021) and was higher in the subsective over privative phrases (*t*(34) = 2.17, *p* = .037).

**Figure 6.**
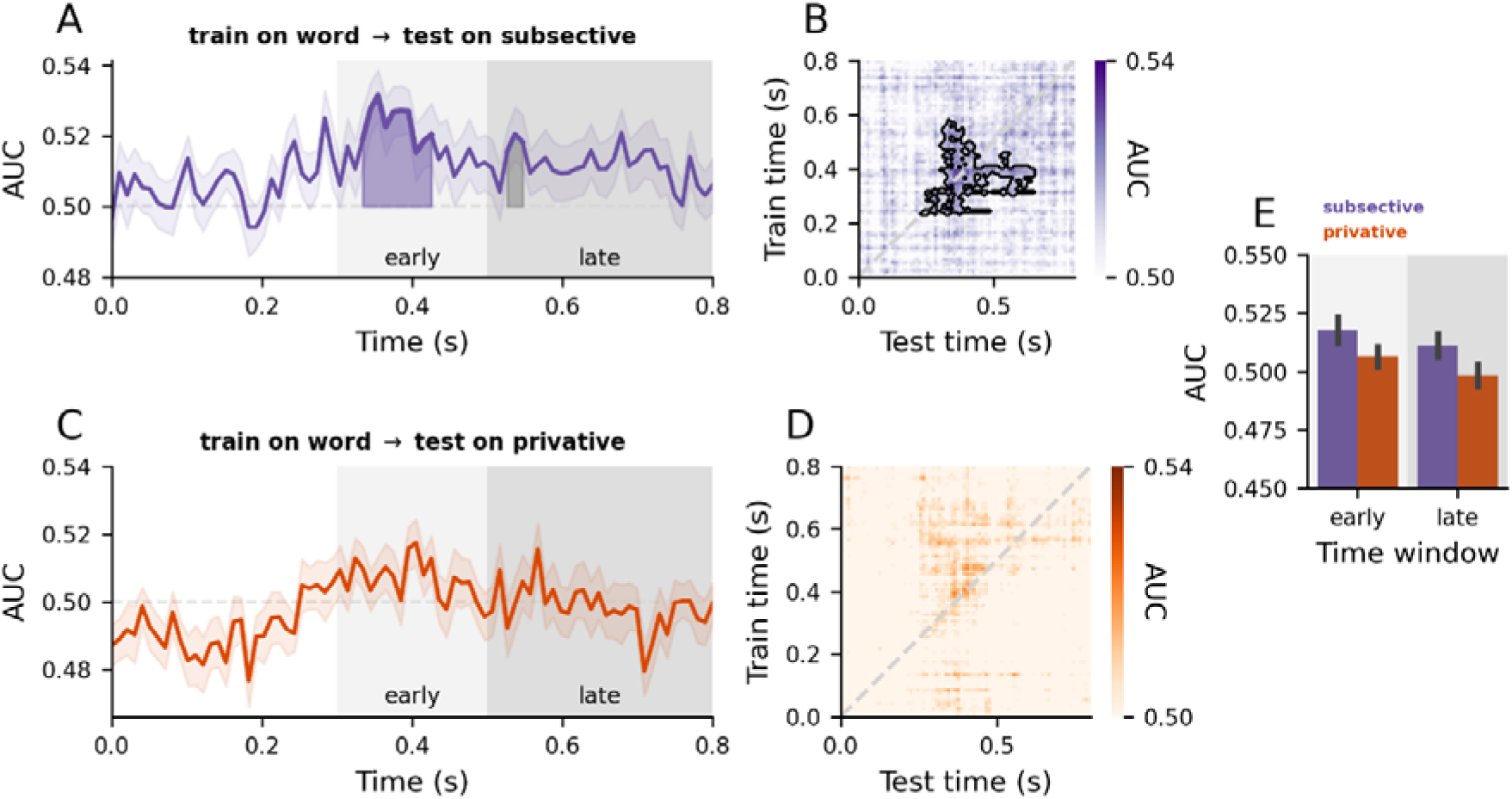
Decoders trained on single word concreteness generalised to subsective but not privative phrases. (A, C) Diagonal decoding performance over time, where time 0 is noun onset. The light and dark grey boxes in the background denote the time window (early vs. late) extents. (B, D) Temporal generalisation matrices, where train time corresponds to single words and test time corresponds to either subsective phrases (B) or privative phrases (D), where time 0 is noun onset. Error bars represent one within-subjects SEM (Loftus & Masson, 1994, 1994). Shaded areas under decoding curves denote clusters corresponding to significant (coloured) and marginally significant (grey) group-level effects assessed with cluster-based one-sample permutation tests against chance. (E) Bar plot showing averaged responses by time window and test type. The light and dark grey boxes in the background correspond to the time window (early vs. late) extents in the time series plots in (A) and (C).

To test the stability of concreteness representation across time, we again performed a temporal generalisation analysis with cluster-based permutation tests, which revealed above-chance decoding in subsective phrases (*p* = .022; Figure 6B) but not in privative phrases (Figure 6D). The cluster had a mean offset from the diagonal of 110 ms and a max offset of 330 ms. The robust cross-condition off-diagonal generalisation suggests that conceptual representations in single words and subsective phrases are relatively consistent in format and that this format is relatively stable in time.

## 4. Discussion

A key challenge in research on semantic cognition, language and their impairments is how meaning is constructed at the level of individual words as well as phrases. These two dimensions of semantics have traditionally been studied separately, within the domains of semantic memory and semantic composition. In integrating these research areas, the present study aimed not only to address fundamental questions about how semantic cognition operates but was also motivated by the observation that both literatures, despite their distinct focuses, consistently highlight the role of the anterior temporal lobe (ATL). In an experimental framework, we investigated whether the activation of conceptual semantic representations from single words and the construction of coherent meaning across word sequences rely on similar neurocomputational mechanisms.

Our analyses revealed that the ATLs are implicated in and sensitive to both concrete and abstract combinatorial semantics across diverse adjective-noun combinations. Bilateral ATLs responded more strongly to phrases than words, independent of concreteness (Figure 3). The left ATL was also sensitive to logical aspects of semantics introduced by subsective vs. privative adjectives (Figure 4). Multivariate decoding analyses on source-localised ATL signals further revealed that the representational dynamics of adjective and noun semantics were distinct: the format of adjective representations dynamically changed over time, while noun representations were relatively more stable in time (Figure 5). By using a cross-decoding regime, we showed that adjectives shape noun semantic representations, with concreteness signals in single words being in a more consistent format to those in subsective phrases than privative phrases (Figure 6).

Given these findings, a parsimonious explanation for the overlap in anatomical and functional localisation appeals to shared computations across semantic memory and semantic composition. We speculate that the ATLs form a representational basis not just for single concepts but also for composed concepts. The system does so by extracting non-linear transmodal and transtemporal statistical structures (Jackson et al., 2021). This form of unified mechanism would shape semantic cognition across ontogenetic and chronometric time scales.

If this notion is correct, we should expect the compositional engines within the ATL to be similarly sensitive to concrete and abstract input concepts. This is exactly what we found. Bilateral ATLs responded more strongly to phrases than words, independent of concreteness, at around 400-650 ms post noun onset (Figure 3). This finding is also compatible with the proposal put forward by Hoffman and colleagues (2018). With a single shared set of “hub” units, their model acquired semantic representations of concrete and abstract concepts as well as their context dependence across sequences of words. More broadly, these insights find echoes in recent advances in theories that argue for a joint role of experiential as well as language-based information in shaping semantic representation (Vigliocco et al., 2009).

Our multivariate decoding analyses provided nascent evidence of how ATL conceptual representations dynamically transform during combinatorial processing. Single-word concreteness decodability persisted in subsective phrases but diminished in privative phrases (Figure 6A & 6C). For subsective adjectives, this suggests a form of non-linear modification where the noun’s conceptual representational format, including its concreteness dimension, is largely preserved. In contrast, the failure to decode concreteness in privative phrases hints at a more profound transformation in the neural representation of conceptual semantics, reflecting the strong non-linear effects privative adjectives induce. Moreover, decodability was stronger in earlier time windows and diminished as processing unfolds (Figure 6E). The temporal generalisation results further underscored this distinction in representational stability. These findings provide evidence that the ATL conceptual representations change in a dynamic, non-linear fashion during combinatorial processing. This capacity for non-linear processing during composition aligns with ATL’s established role in semantic memory (Rogers et al., 2021), where the ATL extracts non-linear statistical structures across modalities and across time to form stable, coherent concepts, thus supporting a unified function.

Our framework offers an explanation for the divergence in effect latencies between our study (400-650 ms) and in prior MEG work targeting the left ATL during combinatorial processing (200-250 ms). The early effect is hypothesized as reflective of “quick-and-easy” (Kim & Pylkkänen, 2019, p. 19) combinations (Poortman & Pylkkänen, 2016; Ziegler & Pylkkänen, 2016; Kim & Pylkkänen, 2019; Pylkkänen & Brennan, 2020). For example, Ziegler and Pylkkänen (2016) reported early effects for adjective-noun phrases whose adjectives were relatively context-insensitive and thus afforded ‘straightforward’ composition. For instance, in the expression “wooden dish”, the adjective ‘wooden’ might contribute a feature [+is-wooden] to the semantic representation of DISH. Whereas, for “large dish”, the adjective ‘large’ encodes/specifies the *relative* size of ‘dish’; this encoding/specification of relative values based on context in scalar adjectives such as ‘large’ or ‘fast’ is thought to be less straightforward compared to adjectives like ‘wooden’. The early effects subsequently diminish for phrases with context-sensitive adjectives which require additional computations. In the present study, conceptual combination demanded by our stimuli is not ‘straightforward’ in this way. Thus, conceptual combination in our case did not manifest in neural readout as an early but later composition effect.

Our subsective-privative contrast revealed a modest univariate effect in the left ATL around 380-440 ms post noun onset. Moreover, cross-decoding analyses from words to phrases revealed better decodability in subsective phrases and at earlier latencies (e.g., 300-500 ms post noun onset) (Figure 6). These data points stand in contrast to prior findings of late (500-800 ms) ERP modulations (Fritz & Baggio, 2020), which was interpreted as reflective of interpretive processes following conceptual access and composition (Baggio, 2018). The apparent divergence in effect timings might be prosaically explained by divergent experimental conditions (e.g., differences in stimulus sets, tasks, or measurements). We believe this is unlikely. Our stimulus set sampled across diverse conceptual domains (including abstract concepts) and contained no repeating items. Despite this increased heterogeneity, which contrasts with the more constrained stimuli typically used, we were still able to detect subtle neural effects, rendering a simple methodological explanation for the timing differences unlikely. Instead, the data appear to support processing models in which the computation of conceptual representations from words and phrases rests on a single, continuous process as opposed to discrete stages. More broadly, our dataset—which includes a contrast targeting logical semantics—contributes to the semantic memory literature in that logical dimensions of meaning should form part of the explanandum, echoing prior work that has begun bridging semantic memory, linguistics, and neurobiology (Fritz & Baggio, 2020, 2022; Westerlund & Pylkkänen, 2014; Zhang & Pylkkänen, 2015, 2018b).

Our analyses revealed right ATL involvement, in addition to the left, during the conceptual processing of phrases, raising interesting questions about the role of the right ATL. The right ATL, like the left ATL, responded more strongly to phrases than words, regardless of phrase concreteness (Figure 3), implying shared processes across the ATLs. Indeed, the proposal that both ATLs are functionally unified during conceptual processing has previously been advanced (Patterson et al., 2007; Schapiro et al., 2013; Lambon Ralph, 2014), with graded left-right asymmetries emerging from differential connectivity between the ATLs and different input/output modality systems (Lambon Ralph et al., 2001; Rice, Hoffman, et al., 2015; Rice, Lambon Ralph, et al., 2015; Schapiro et al., 2013). For example, linguistic stimuli would engage the left ATL to a greater extent. Indeed, this pattern of results was borne out in our dataset (Figure 3B). Together, the results obtained here suggest that semantic memory and composition are unified both within *and across* the ATLs, complementing prior hemodynamic work and supporting an account of a bilateral-yet-graded, connectivity-constrained ‘hub’ for semantic cognition.

The present study also paves the way for models of semantic cognition to better understand time-extended semantic cognition beyond simple adjective-noun phrases. Incorporating phrasal-level manipulations into models of semantic cognition could broaden their empirical coverage from single words through to time-extended coherent verbal narratives. A straightforward next step would be to extend the present approach to explore the conceptual processing of other linguistic elements, such as adjectives and verbs. For example, neuropsychological evidence from SD patients suggests that ATL similarly provides a representational basis for event concepts (denoted by verbs) and entity concepts (denoted by nouns) alike (Patterson et al., 2001; Bonner et al., 2009). MEG investigations on the composition of verb phrases also implicated the ATL (Kim & Pylkkänen, 2019). The approach taken in this study is thus well-positioned for further linking between semantic memory and composition. Our results have shown that adjective and noun semantics were represented with distinct dynamics (Figure 5B & D). Future work could explore computational models of semantic representation of different phrase types (e.g., noun vs. verb phrases) to explain the representational differences observed in our study. Moreover, model-internal representations and neural representations in humans could be linked for a potential mechanistic understanding of semantic cognition. Indeed, this is an approach that is commonplace in vision neuroscience (e.g., Cichy et al., 2014; Kietzmann et al., 2019) and has recently been applied to studies of semantic cognition (Rogers et al., 2021).

Ultimately, we construct meaning out of rich, multimodal communicative situations and contexts. Such a unified framework for representing and integrating conceptual knowledge over time paves the way for new and exciting research questions about the nature of *natural* semantic cognition. For example, when communicating, we often use language together with non-linguistic cues, such as gestures (e.g., when giving directions, describing the size of an object). The conceptual processing of co-speech gestures has been extensively studied in brain and behaviour (for a review of general non-linguistic conceptual processing informed by the N400, see e.g. Kutas & Federmeier, 2011; for review of co-speech gesture processing, see e.g. Özyürek, 2014). Further, semantic composition in American Sign Language has also been shown to engage the left ATL (Blanco-Elorrieta et al., 2018). Thus, future work could explore the neurocomputational bases of the integration of conceptual information from different linguistic and non-linguistic sources.

## Data and Code Availability

The data will be made available at http://www.mrc-cbu.cam.ac.uk/publications/open444data. Code and analysis pipelines will be made available at https://github.com/rmc-law/FakeDiamond.

## Author Contributions

R.M.C.L.: Conceptualisation, Methodology, Software, Validation, Formal Analysis, Investigation, Writing – Original Draft, Writing – Review & Editing, Visualization, Project Administration, Data Curation

O.H.: Conceptualisation, Methodology, Resources, Writing – Review & Editing, Supervision, Project Administration, Funding Acquisition

M.A.L.R: Conceptualisation, Methodology, Resources, Supervision, Writing – Review & Editing, Project Administration, Funding Acquisition

All authors reviewed and approved the final manuscript.

## Acknowledgements

We are grateful to Clare Cook, Lucy MacGregor, Steve Eldridge, Máté Aller, Rebecca Williams, Fatih Serin, Marius Mada, and Karen Kabakulu for their assistance with data collection and also to Tobias Goehring and Lidea Shahidi for their helpful advice on data analysis. We are also grateful to Hugo Weissbart for initial guidance on decoding analyses for a separate project.

## Author notes

Open access: For the purpose of open access, the UKRI-funded authors O.H. and M.A.L.R. have applied a Creative Commons Attribution (CC BY) license to any Author Accepted Manuscript version arising from this submission.

## Supplementary Material

**Supplementary Figure 1.**
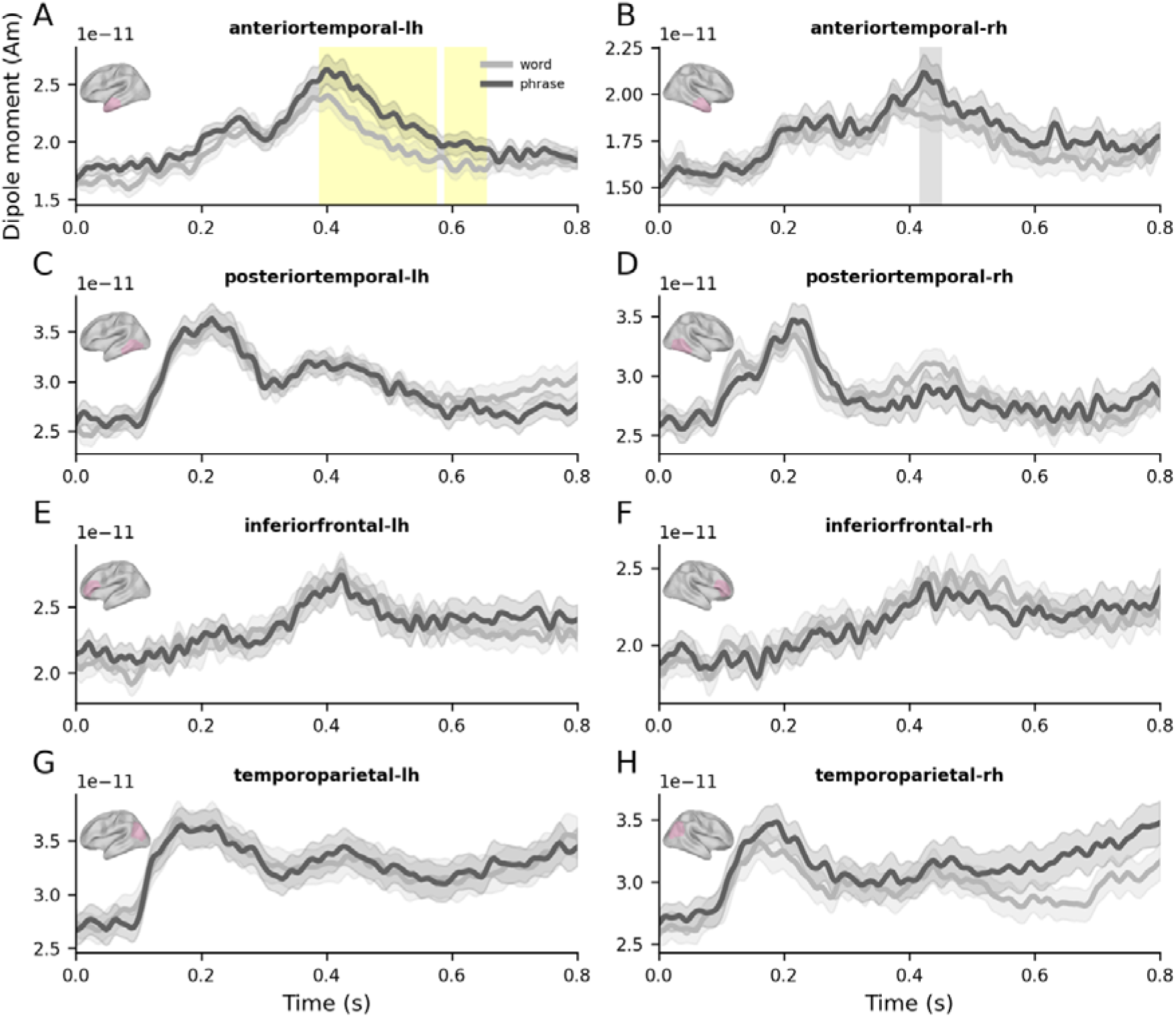
Composition effects in each tested ROI. (A-H) Time series of source-localised evoked responses for words and subsective phrases, aligned to noun onset (0 s). Error bars represent ±1 within-subjects SEM (Loftus & Masson, 1994). Cluster extent identified from permutation testing are indicated by yellow shaded areas for significant effects (p<.05) and grey shaded areas for marginally significant effects (p<.1). Of the detectable effects across ROIs, effects in the left ATL survived FDR correction across multiple ROI comparisons (q = .006).

**Supplementary Figure 2.**
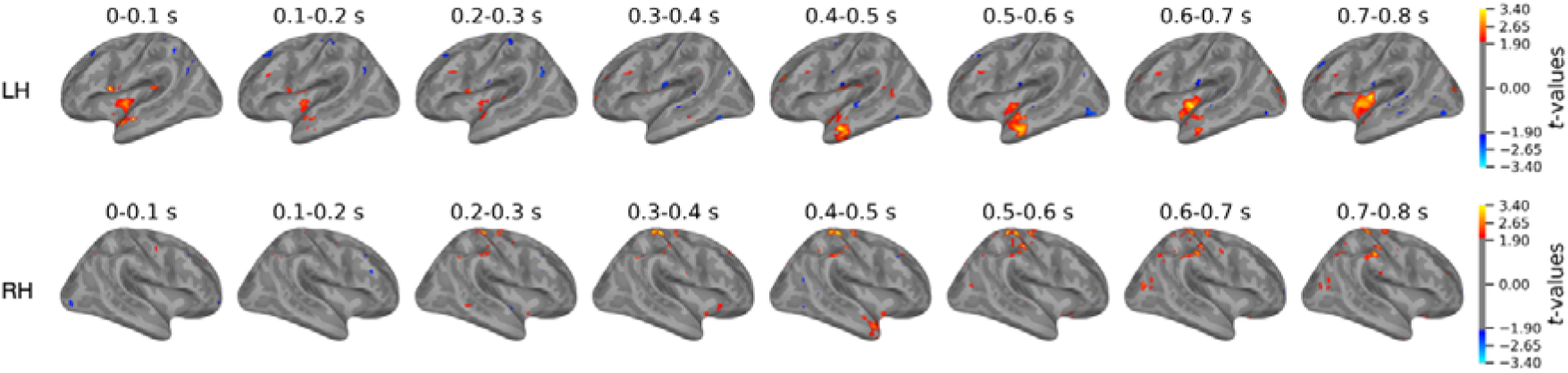
Spatiotemporal (uncorrected) t-maps of composition effects (phrase word) across both the left and right hemispheres. Each brain model depicts t-maps binned with each 100 ms wide time window.

**Supplementary Figure 3.**
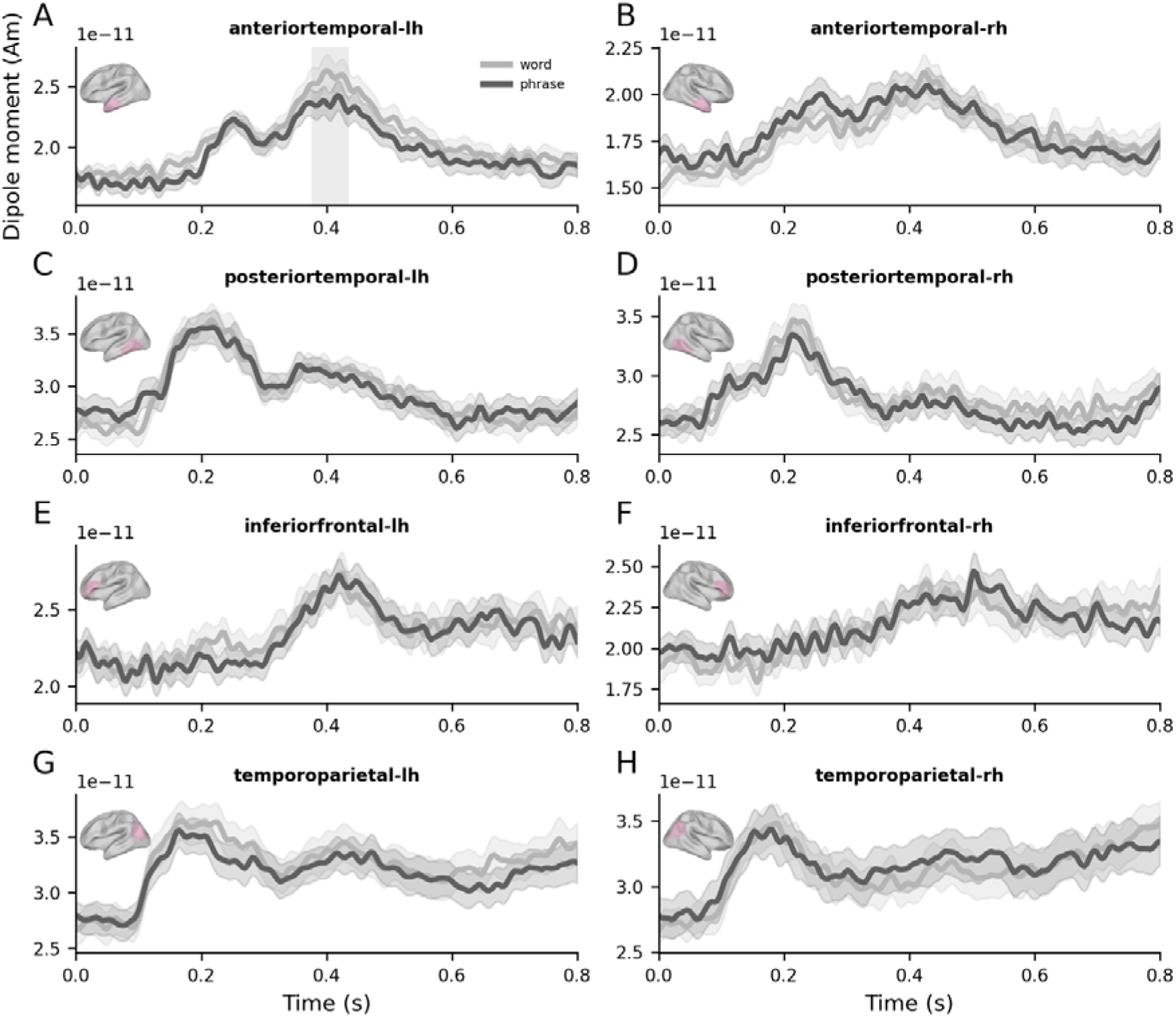
Denotation effects in each tested ROI. (A-H) Time series of source-localised evoked responses for subsective and privative phrases, aligned to noun onset (0 s). Error bars represent ±1 within-subjects SEM (Loftus & Masson, 1994). Cluster extent identified from permutation testing are indicated by yellow shaded areas for significant effects (p<.05) and grey shaded areas for marginally significant effects (p<.1).

**Supplementary Figure 4.**
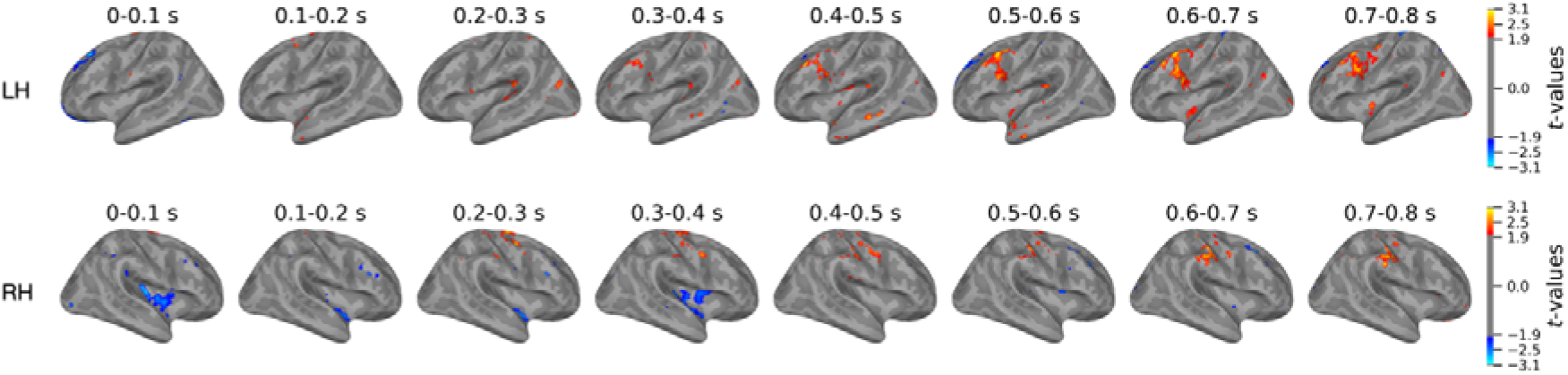
Spatiotemporal (uncorrected) t-maps of denotation effects (subsective privative) across both the left and right hemispheres. Each brain model depicts t-maps binned with each 100 ms wide time window.

**Supplementary Figure 5.**
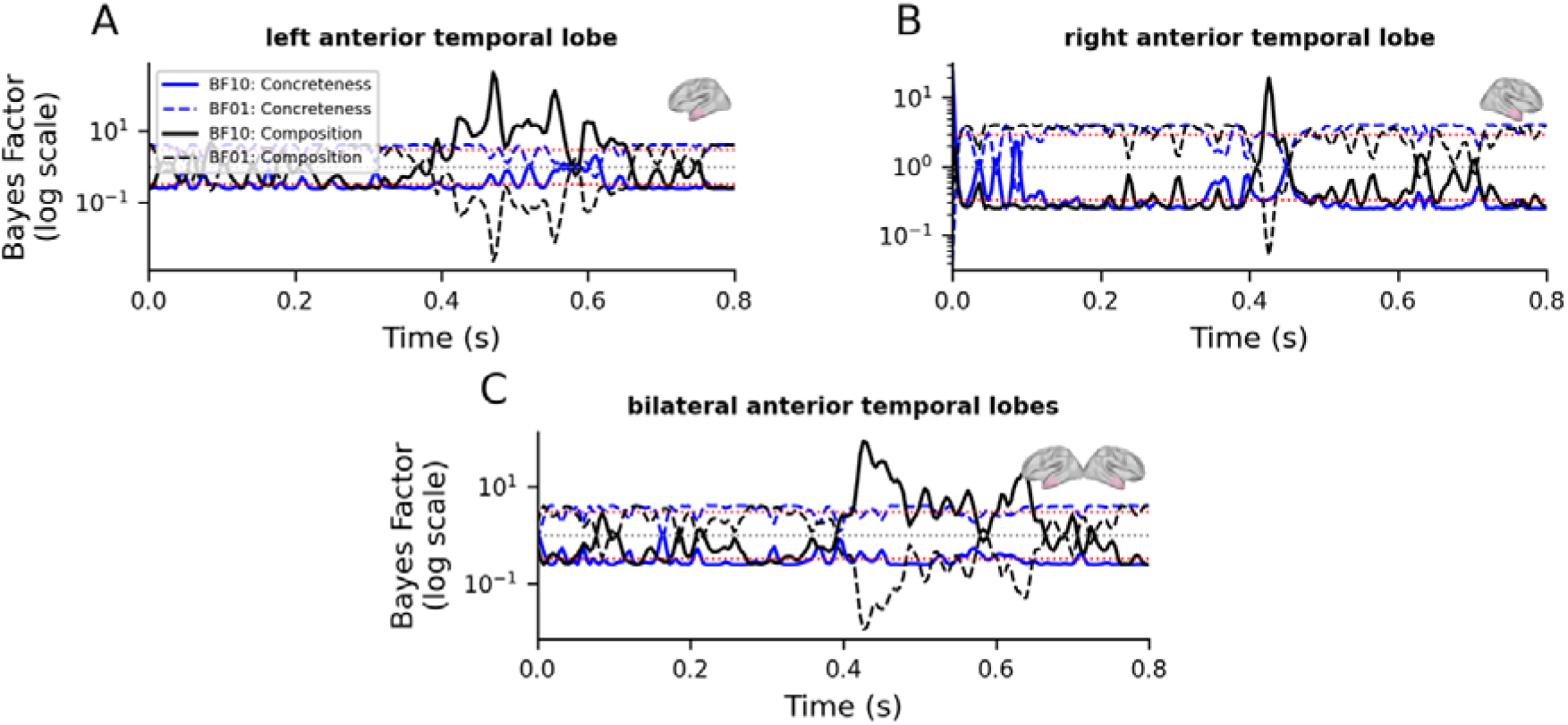
Time series of Bayes Factor illustrating the strength of evidence for effects of Composition (black solid lines), Concreteness (blue solid lines) over time. The corresponding evidence for the null hypothesis (BF01) is shown with dashed lines of the same colour. The Bayes Factor is plotted on a logarithmic scale (y-axis). Horizontal dotted lines indicate thresholds for evidence: BF=1 (grey dotted line) indicates no evidence for either the alternative or the null hypothesis; BF=3 and BF=1/3 (red dotted lines) indicate moderate evidence for the alternative and null hypotheses, respectively. Each subplot includes an inset brain image indicating the anatomical location of the anterior temporal region of interest (ROI).

**Supplementary Figure 6.**
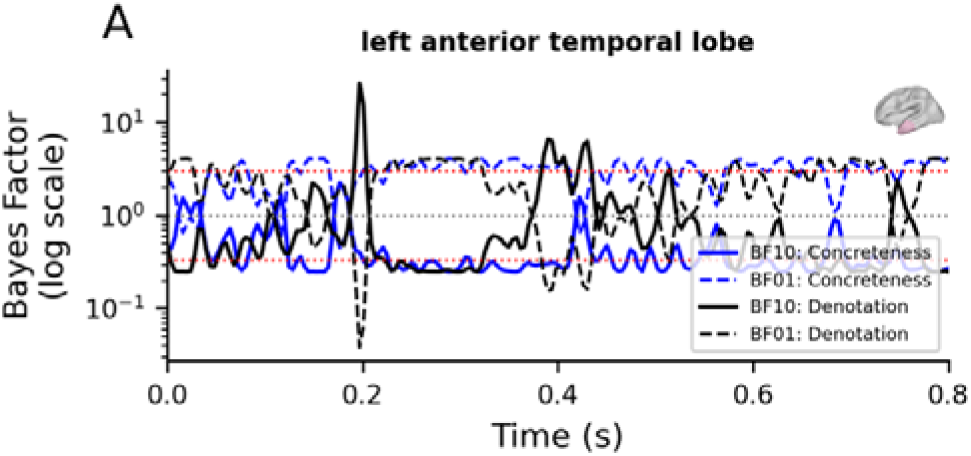
Time series of Bayes Factor illustrating the strength of evidence for effects of Denotation (black solid line: BF10 in A) over time. The corresponding evidence for the null hypothesis (BF01) is shown with dashed lines of the same colour. The Bayes Factor is plotted on a logarithmic scale (y-axis). Horizontal dotted lines indicate thresholds for evidence: BF=1 (grey dotted line) indicates no evidence for either the alternative or the null hypothesis; BF=3 and BF=1/3 (red dotted lines) indicate moderate evidence for the alternative and null hypotheses, respectively. Each subplot includes an inset brain image indicating the anatomical location of the anterior temporal region of interest (ROI).

