## Supplemental figures for "A common framework for semantic memory and semantic composition"

### Supplementary figures

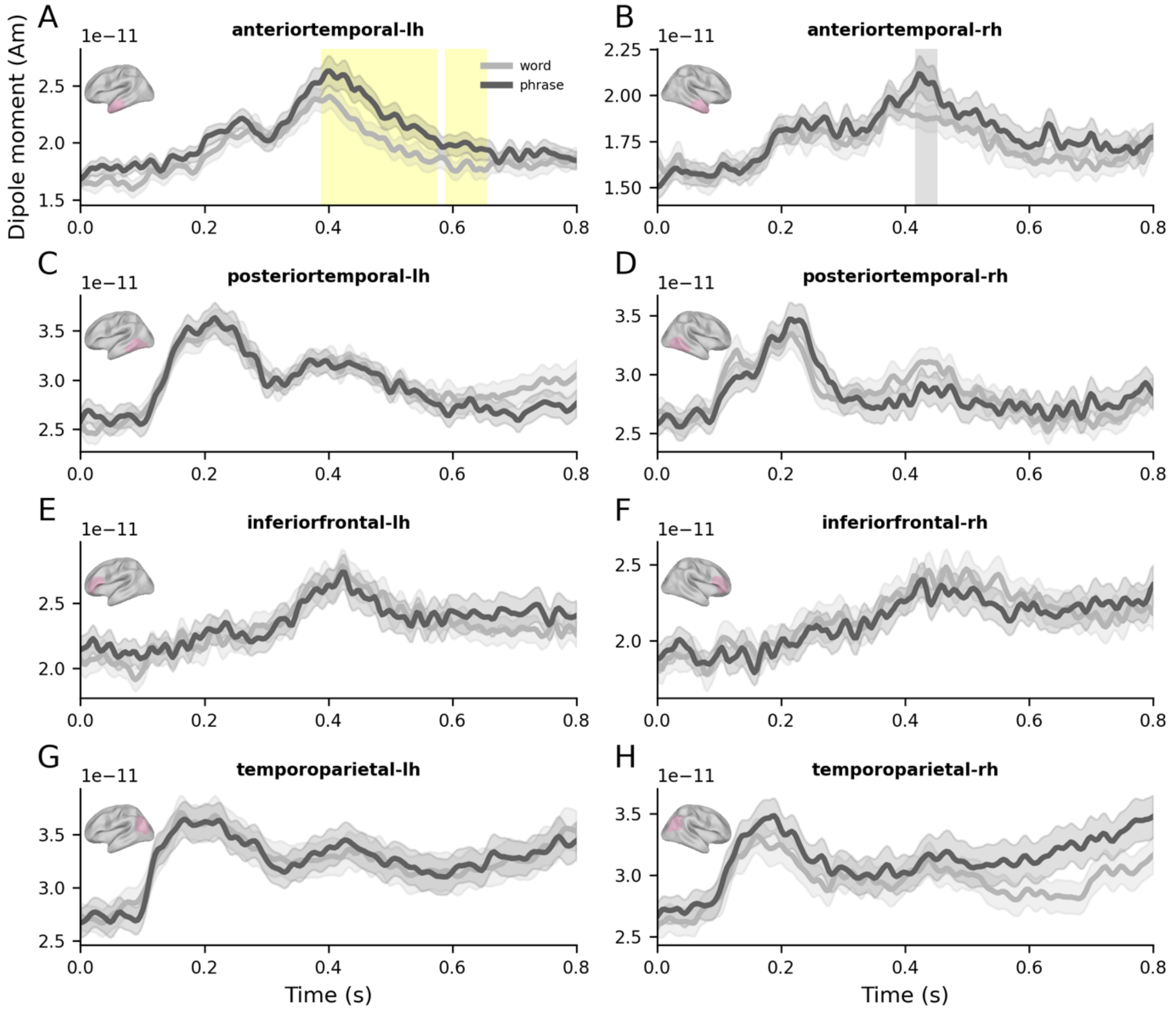


Supplementary Figure *1. Composition effects in each tested ROI. (A-H) Time series of source-localized evoked responses for words and subsective phrases, aligned to noun onset (0 s). Error bars represent ±1 within-subjects SEM (Loftus & Masson, 1994). Cluster extent identified from permutation testing are indicated by yellow shaded areas for significant effects (p<.05) and grey shaded areas for marginally significant effects (p<.1). Of the detectable effects across ROIs, effects in the left ATL survived FDR correction across multiple ROI comparisons (q = .006).*

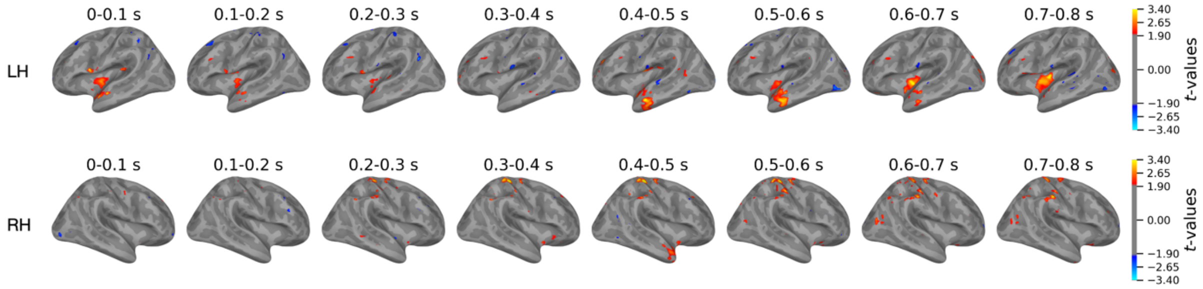


*Supplementary Figure 2. Spatiotemporal (uncorrected) t-maps of composition effects (phrase – word) across both the left and right hemispheres. Each brain model depicts t-maps binned with each 100 ms wide time window.*

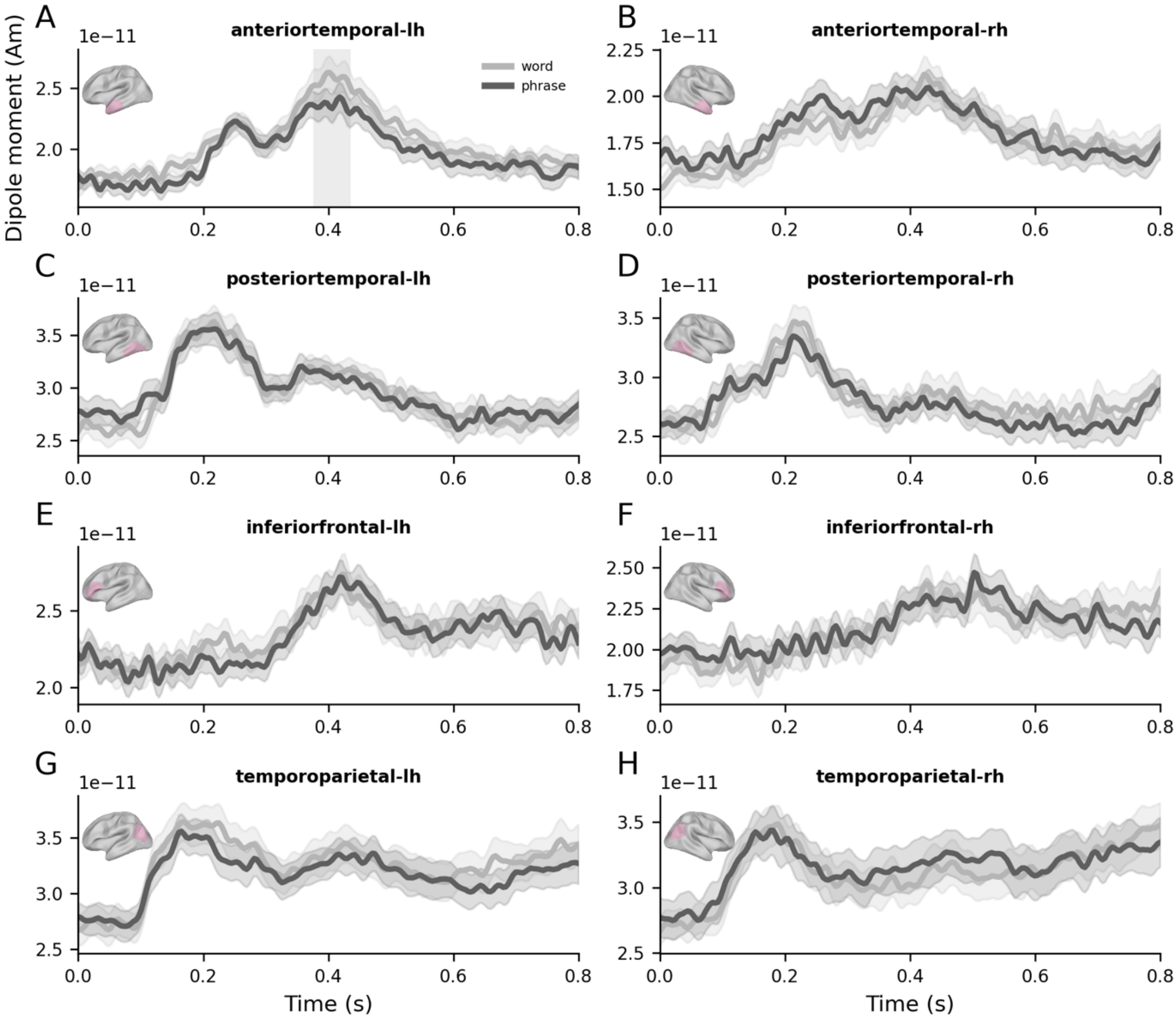


Supplementary Figure *3. Denotation effects in each tested ROI. (A-H) Time series of source-localized evoked responses for subsective and privative phrases, aligned to noun onset (0 s). Error bars represent ±1 within-subjects SEM (Loftus & Masson, 1994). Cluster extent identified from permutation testing are indicated by yellow shaded areas for significant effects (p<.05) and grey shaded areas for marginally significant effects (p<.1).*

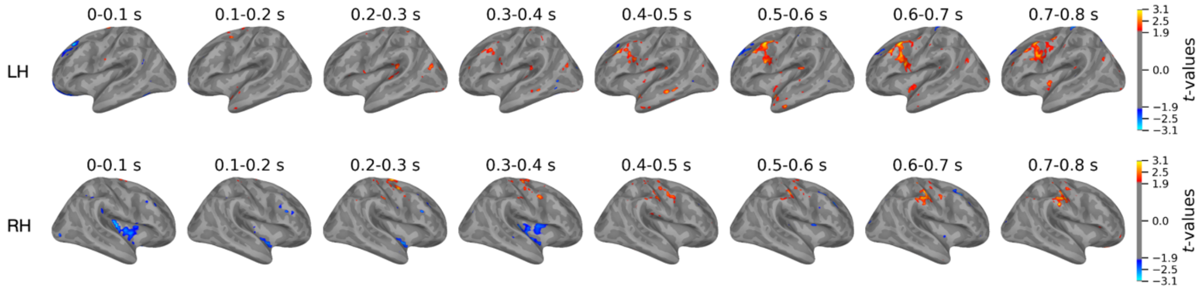


*Supplementary Figure 4. Spatiotemporal (uncorrected) t-maps of denotation effects (subsective – privative) across both the left and right hemispheres. Each brain model depicts t-maps binned with each 100 ms wide time window.*

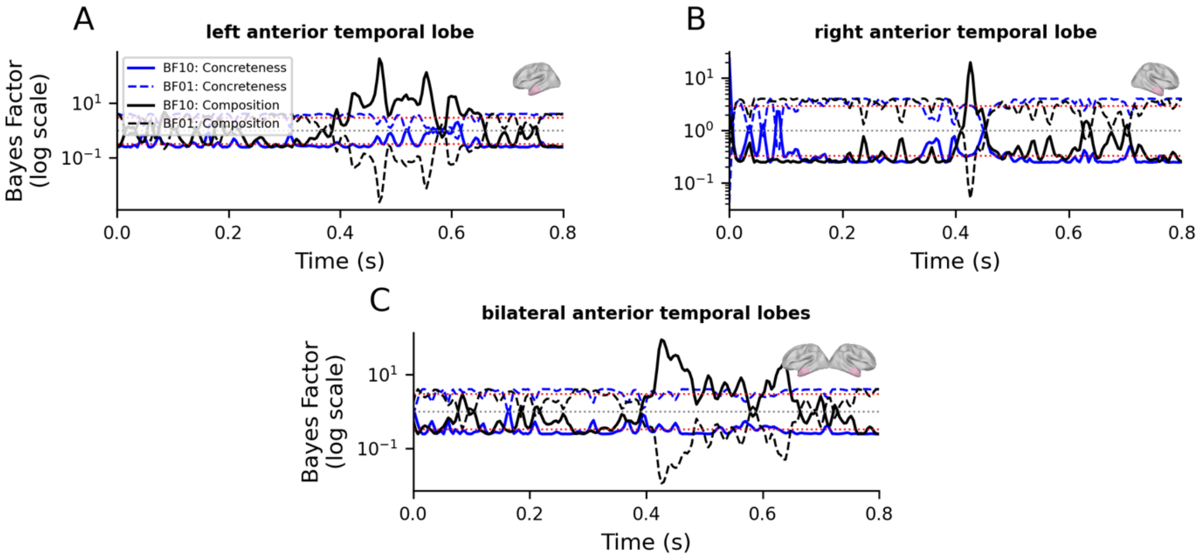


*Supplementary Figure 5. Time series of Bayes Factor illustrating the strength of evidence for effects of Composition (black solid lines), Concreteness (blue solid lines) over time. The corresponding evidence for the null hypothesis (BF01) is shown with dashed lines of the same colour. The Bayes Factor is plotted on a logarithmic scale (y-axis). Horizontal dotted lines indicate thresholds for evidence: BF=1 (grey dotted line) indicates no evidence for either the alternative or the null hypothesis; BF=3 and BF=1/3 (red dotted lines) indicate moderate evidence for the alternative and null hypotheses, respectively. Each subplot includes an inset brain image indicating the anatomical location of the anterior temporal region of interest (ROI).*

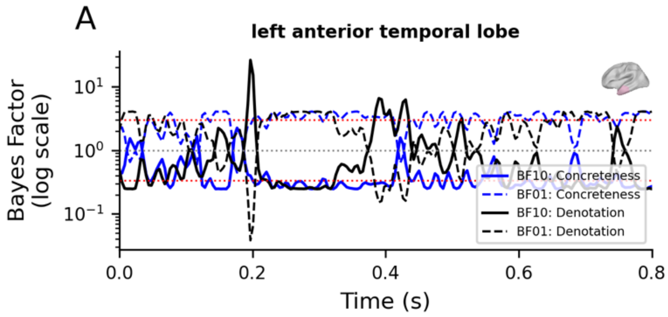


*Supplementary Figure 6. Time series of Bayes Factor illustrating the strength of evidence for effects of Denotation (black solid line: BF10 in A) over time. The corresponding evidence for the null hypothesis (BF01) is shown with dashed lines of the same colour. The Bayes Factor is plotted on a logarithmic scale (y-axis). Horizontal dotted lines indicate thresholds for evidence: BF=1 (grey dotted line) indicates no evidence for either the alternative or the null hypothesis; BF=3 and BF=1/3 (red dotted lines) indicate moderate evidence for the alternative and null hypotheses, respectively. Each subplot includes an inset brain image indicating the anatomical location of the anterior temporal region of interest (ROI).*
